# Complex and long-range linkage disequilibrium and its relationship with QTL for Marek’s Disease resistance in chicken populations

**DOI:** 10.1101/2021.05.27.445970

**Authors:** Ehud Lipkin, Janet E. Fulton, Jacqueline Smith, David W. Burt, Morris Soller

## Abstract

Chicken long-range linkage disequilibrium (LRLD) and LD blocks, and their relationship with previously described Marek’s Disease (MD) quantitative trait loci regions (QTLRs), were studied in an F_6_ population from a full-sib advanced intercross line (FSAIL), and in eight commercial pure layer lines. Genome wide LRLD was studied in the F_6_ population by random samples of non-syntenic and syntenic marker pairs genotyped by Affymetrix HD 600K SNP array. To illustrate the relationship with QTLRs, LRLD and LD blocks in and between the MD QTLRs were studied by all possible marker pairs of all array markers in the QTLRs, using the same F_6_ QTLR genotypes and genotypes of the QTLR elements’ markers in the eight lines used in the MD mapping study. LRLD was defined as r^2^ ≥ 0.7 over a distance ≥ 1 Mb, and 1.5% of all syntenic marker pairs were classified as LRLD. Complex fragmented and interdigitated LD blocks were found, over distances ranging from a few hundred to a few million bases. Vast high, long-range, and complex LD was found between two of the MD QTLRs. Cross QTLRs STRING networks and gene interactions suggested possible origins of this exceptional QTLRs’ LD. Thus, causative mutations can be located at a much larger distance from a significant marker than previously appreciated. LRLD range and LD block complexity may be used to identify mapping errors, and should be accounted for while interpreting genetic mapping studies. All sites with high LD with a significant marker should be considered as candidate for the causative mutation.

## INTRODUCTION

Linkage disequilibrium (LD) refers to correlations among alleles of different genomic sites. It quantifies the informativity between different sites [Reich et al., 2001; Ardlie et al., 2002; Aerts et al., 2007]. Useful LD indicate non-random association of alleles at different loci [Aerts et al., 2007]. Appreciable LD is commonly found between pairs of loci close to one another, and LD decreases rapidly as distance between the loci increases [e.g., Aerts et al., 2007; García-Gámez et al., 2012]. This provides the basis for genome wide association studies (GWAS) to map quantitative trait loci (QTL).

Nevertheless, during any one-time snapshot of a population, long-range LD (LRLD) can also be found among loci that are well-separated from one another, over millions of bp [Aerts et al., 2007; Corbin et al., 2010; García-Gámez et al., 2012; Koch et al., 2013; Skelly et al, 2015; Park 2019; Peters et al., 2021]. Rarely, LD above background level can be found between non-syntenic markers on different chromosomes (chr) [García-Gámez et al., 2012].

LRLD could just be a matter of sampling variation, especially in small populations [Skelly et al, 2015]. Alternatively, LRLD could be a result of genome assembly errors where SNP locations are misidentified, and thus LRLD may help identify such assembly errors [Utsunomiya et al., 2016]. This phenomenon may, however, also have genuine biological origins, such as co-evolution of genomic sites (coding and non-coding genes, long- and short-range regulatory sites), gene conversion, copy number variation, or demographic factors such as selection, population bottlenecks, nonrandom mating, and epistasis.

LD blocks are runs of genomic sites all having appreciable LD with one another. However, high LD in general and LRLD in particular are not always continuous. Rather, they are often separated by genomic sites with which they have practically no LD [Allabi et al., 2005; Aerts et al., 2007; Lipkin et al., 2013; O’Brien et al., 2014]. Thus, not only can LD be found among distant loci, but also its pattern may be complex, comprised of fragmented blocks.

LRLD and LD complexity present concerns for GWAS mapping, as a significant association nay be found between a causative locus and markers far removed from it, thus falsely placing the putative causative locus at a site far away from its actual location [Skelly et al, 2015]. On the other hand, LRLD may point to interactions between unlinked regions in the genome (e.g., a receptor and its ligand or a gene and its regulator). Furthermore, LRLD can identify co-evolution of different genomic regions affected by the same selection, natural or artificial.

The objectives of this study were to characterize LRLD and LD blocks in multiple Hy-Line chicken lines previously used to map quantitative trait loci regions (QTLRs) for MD resistance [Smith et al., 2020]. This will give a genomic view of the LD complexity, and illustrate its importance to mapping results by assessing the relationship between LRLD and LD blocks and MD QTLRs.

## MATERIALS AND METHODS

### Populations

All procedures carried out on the birds involved in this study were conducted in compliance with Hy-Line International Institutional Animal Care and Use Committee guidelines.

Nine populations described by Smith et al. (2020) were again used in the present study. These comprised five families of F_6_ birds from a Full Sib Advanced Intercross Line (FSAIL) used by Smith et al. (2020) to map QTLRs affecting MD resistance, and eight pure lines used in the same study to test these QTLRs.

A priory, it is expected that LRLD in F_6_ will be at higher frequency than in pure lines, because the families start with only 4 chromosomes each, and there are only a few generation of intercrossing to break up the haplotype blocks. However, this study did not compare populations, families or lines. Rather, it aim to present the phenomena of LD long range and complexity.

### Trait

The trait for which these QTLR are associated is resistance to the avian oncogenic alpha herpes virus, Marek’s Disease (MD virus [Smith et al., 2020]. This trait association data set was used, as the phenotype and genotype information was available. It is used as an illustration for the QTLR and LD associations that were identified.

### Genotypes

Only the autosomal genotypes as used by Smith et al. (2020) to map and test QTLRs were used in the present study. Genotypes on the Z chromosome will be analyzed in detail in a different manuscript. Genotypes were obtained from the HD 600K Affymetrix SNP chicken array [Kranis et al., 2013] in the F_6_ population, and marker genotypes used to test the QTLRs by the eight pure lines [Smith et al., 2020]. The only difference was that instead of a minimum MAF ≥ 0.01 used for the association tests by Smith et al. (2020), a threshold of 0.10 was used here for the LD analysis, to avoid spurious high LD due to rare alleles [Skelly et al., 2015]. However, to test LD with exactly the same markers used for association tests in the eight pure lines, the threshold of MAF ≥ 0.01 was also used for LD for these lines.

### Genome assemblies and remapping QTLRs

As described in Smith et al. (2020), the initial analysis in this study was based on the Galgal4 genome build. The <u>Lift Genome Annotations</u> tool [Haeussler et al., 2019] within the <u>UCSC</u> Browser was used to remap markers from Galgal4 to Galgal6 (GRCg6a; Acc. No.: GCA_000002315.5). The new coordinates were then used to remap the F_6_ QTLRs as was done by Smith et al. (2020). The results were very similar on both assemblies, and hence only the results from Galgal6 will be presented here. Nevertheless, there was one change worth noting, detailed in the Appendix.

### Linkage disequilibrium (LD)

#### LD Measure

LD r^2^ within each F_6_ family and each pure line were obtained using JMP Genomics software (JMP Genomics, Version 9, SAS Institute Inc., Cary, NC, USA, 1989–2019).

#### Non-syntenic LD

Background LD over all autosomes was estimated utilizing Affymetrix 600K genotypes of two random combined samples with return of non-syntenic marker pairs from each of the five F_6_ families.

#### Long-Range LD (LRLD)

Koch et al. (2013) used all pairs of SNPs on each human chromosome and defined LRLD between haplotype-blocks rather than between SNP pairs. However, due to computer limitations, we used samples of random SNP pairs on the Affymetrix 600K SNP array to assess LRLD in the autosomes of the five F_6_ families. Koch et al. (2013) defined LRLD in human as high LD (their low *p*_D_) over a distance ≥ 0.25 cM (≈ 0.25 Mb). Vallejo et al. (2018) used a very relaxed LD threshold of r^2^ > 0.25 in rainbow trout studies, but a larger minimum distance of 1.0 Mb. Conservatively, and based on the present results, in this study we defined LRLD as a marker pair with r^2^ ≥ 0.7 over a distance ≥ 1.0 Mb. As noted in the Introduction, LRLD between elements in a QTLR and across QTLRs can indicate a relationship between the element and between the QTLRs. Hence, to illustrate its importance, LRLD in and between MD QTLRs [Smith we al., 2020] was studied by all F_6_ Affymetrix genotypes in the QTLRs.

#### F_6_ random LRLD and QTLRs

F_6_ LRLD of random marker pairs were aligned by chromosomal location with the F_6_ MD QTLRs [Smith et al., 2020], and all overlaps between the LRLDs and the QTLRs were counted.

#### LD blocks

High LD blocks were defined as a group of markers located on the same chromosome having r^2^ ≥ 0.7 with each other. The definition was applied even if markers with low LD appeared between the LD markers. This definition allowed a “look over the horizon” and identification of fragmented and interdigitated blocks.

### Gene network analysis

To investigate the genes underlying QTLRs 4 and 5, the BioMart tool within the Ensembl database (https://www.ensembl.org/info/data/biomart/index.html) was used to identify genes in these regions. These identified genes were then subject to network analysis using the STRING database (v11). [Jensen et al., 2009] which provides an overview of known protein interactions.

## RESULTS

### Remapping QTLRs from Galgal4 to Galgal6

The new coordinates of the markers on Galgal6 and association results obtained by Smith et al. (2020) were used to remap the MD QTLRs identified with Galgal4 by Smith et al. (2020). The same 38 QTLRs were found on each genome build (Table 1). Most changes were negligible, except one movement of a fragment on Chr 1 over about 70 Mb from QTLR 1 to QTLR 4, including a QTLR lncRNA tested by Smith et al. (2020). This change is detailed in the Appendix. The new QTLR coordinates on Galgal6 were used for the LD analyses in this study.

**Table 1.**
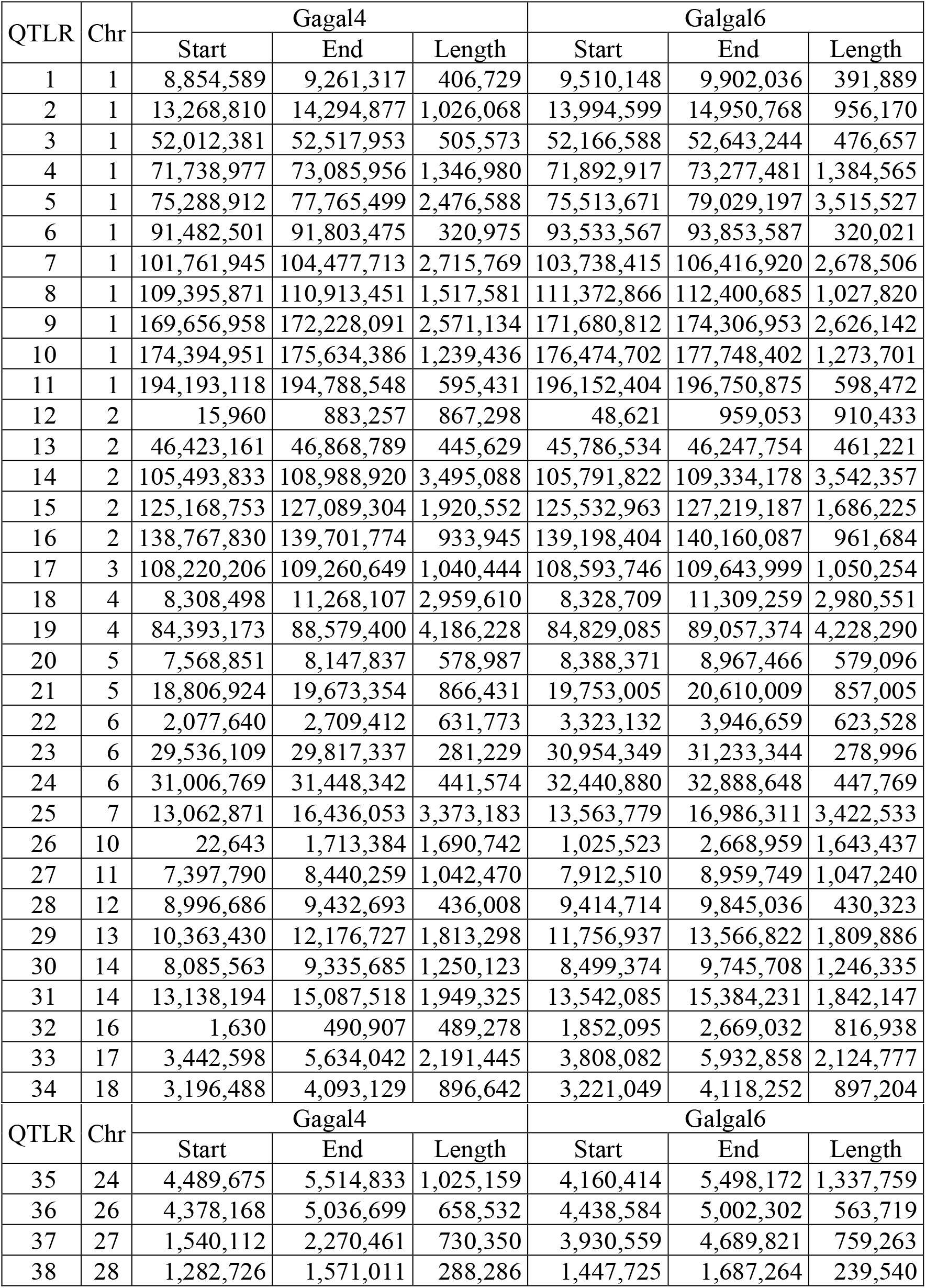
Remapping Galgal4 QTLRs onto Galgal6. QTLR, ordinal number of the QTLR [Smith et al., 2020]; Chr, chromosome; Start/End, QTLR coordinates of the first and last SNP in the QTLR; Length, size of the QTLR (bp).

### Linkage disequilibrium in the F_6_ population based on 600K genotyping SNP array

#### Non-syntenic random LD

A total of 923,183 random non-syntenic pairs of markers from different autosomes were used to assess the background level of LD, with an average of 184,636.6 pairs in a family (Table 2). LD averaged 0.011 ± 0.016, comparable to previous reports in chicken [Lipkin et al., 2013; Seo et al., 2018], but about ten times higher than values reported in mammals; horse [Corbin et al., 2010), sheep [García-Gámez et al., 2012], and cattle [Khatkar et al., 2008; Sargolzaei et al., 2008] populations. These differences may represent experimental design, or population sample, size, history and structure or biological differences between birds and mammals.

**Table 2.** Summary statistics of non-syntenic random LD between 600K markers in the F_6_ families. Pairs, number of pairs r^2^ values obtained; Ind/pairs, average number of individuals used to calculate a markers pair LD in a family; r^2^: Avg, average; SD, standard deviation; Min, minimum; Max, maximum; All: Pairs, average number of pairs in all families combined; Ind/pairs r^2^, means weighted by the number of Pairs.

| Family | Pairs | Ind/<br>pair | r<sup>2</sup> |  |  |  |
| --- | --- | --- | --- | --- | --- | --- |
|  |  |  | Avg | SD | Min | Max |
| 1 | 190,232 | 176.6 | 0.014 | 0.019 | 0.000 | 0.368 |
| 2 | 182,366 | 231.7 | 0.012 | 0.016 | 0.000 | 0.991 |
| 3 | 184,399 | 354.0 | 0.008 | 0.011 | 0.000 | 0.372 |
| 4 | 187,406 | 219.5 | 0.010 | 0.014 | 0.000 | 0.246 |
| 5 | 178,780 | 200.7 | 0.012 | 0.016 | 0.000 | 0.243 |
| All | 184,636.6 | 236.3 | 0.011 | 0.016 | 0.000 | 0.443 |

Mean family LDs and standard deviations (SDs) had significant high negative correlation with the size of the population sample (i.e., Ind/pair), r = −0.895 (P = 0.040) and r = −0.884 (P = 0.046). These correlations are also in accord with our previous report in chicken [Lipkin et al., 2013], and the expectation of Sved (1971).

With a mean LD of 0.011 ± 0.016, any r^2^ > 0.043 is above the background LD. Indeed, combining all five F_6_ families together, only 3.5% of the r^2^ values were above 0.05 (Supplemental Table 1). A single high LD of r^2^= 0.991 was found in Family 2. Without any replication, this was treated as a sampling effect. Based on these results, a conservative critical LD value of r^2^ ≥ 0.15 was chosen for defining significant LD.

***Supplemental Table 1**. Distribution of non-syntenic random LD values among the F_6_ families*.

#### Syntenic random LD

A total of 1,008,823 random syntenic marker pairs were used to assess the level of random LDs on the same chromosome (Table 3). Distance between markers in the random pairs varied from 11 to 197,038,449 bp, with an average of 28,976,195.6 bp. Means of r^2^ were all in close range around 0.11, averaging 0.114, ten times the means obtained for the non-syntenic LD. Though obtained by random marker pairs, some of which are at a long distance from one another, these means suggest the presence of large number of LDs above the background LD of 0.011 (Table 2). The expected negative correlation between distance and LD was again obtained in all five families (Table 3).

**Table 3.** Summary statistics of syntenic random LD between 600K markers in the F_6_families. Pairs: No., number of pairs r^2^ values obtained; Ind, average number of individuals used to calculate a markers pair LD in a family; bp: average, minimum and maximum bp between markers in a pair; r^2^: Avg, average; SD, standard deviation; Min, minimum; Max, maximum; r(bp-r^2^), correlation between the distance and r^2^ of a marker pair; All: Pairs, total number of pairs in all families combines; Ind, bp and r^2^: means of weighted by the number of pairs.

| Family: |  | 1 | 2 | 3 | 4 | 5 | All |
| --- | --- | --- | --- | --- | --- | --- | --- |
| Pairs | No. | 207,770 | 200,411 | 200,866 | 204,453 | 195,323 | 1,008,823 |
|  | Ind | 176.6 | 231.7 | 354.0 | 219.5 | 200.7 | 236.2 |
| bp | Avg | 28,745,980.9 | 30,164,000.4 | 28,092,640.5 | 29,039,387.5 | 28,844,817.9 | 28,976,195.6 |
|  | Min | 11 | 11 | 40 | 64 | 165 | 57.3 |
|  | Max | 196,356,451 | 197,038,449 | 196,167,202 | 196,407,037 | 196,676,774 | 196,526,525.6 |
| $r^2$ | Avg | 0.119 | 0.112 | 0.109 | 0.110 | 0.119 | 0.114 |
|  | SD | 0.221 | 0.218 | 0.216 | 0.218 | 0.224 | 0.220 |
|  | Min | 0.000 | 0.000 | 0.000 | 0.000 | 0.000 | 0.000 |
|  | Max | 1.000 | 1.000 | 1.000 | 1.000 | 1.000 | 1.000 |
| $r(\text{bp}-r^2)$ | | -0.346 | -0.346 | -0.347 | -0.336 | -0.351 | -0.345 |

In all families, about two-thirds of the LD values were up to 0.05, dropping rapidly thereafter (Table 4). Interesting, for all families, there was an increase in the range of r^2^> 0.85, suggesting existence of large high-LD blocks. Pooled over all families, the proportion of r^2^ ≥ 0.15, set conservatively as a threshold of significance by the non-syntenic LD, was almost 0.2 (Table 4), while less than 5% of the LD values were above 0.7. Hence, the range of 0.15 ≤ r^2^ < 0.7 was set as low to moderate LD and used to define moderate LD blocks, and r^2^ ≥ 0.7 was set as high LD and used to define LRLD and high LD blocks.

**Table 4.** Distribution of syntenic random LD values among the F_6_ families. All, all families combine; r^2^ ≥ (last 2 rows), frequencies of r^2^ above the indicated value.

| $r^2$ | Fam1 | Fam2 | Fam3 | Fam4 | Fam5 | All |
| --- | --- | --- | --- | --- | --- | --- |
| $\leq 0.05$ | 0.6429 | 0.6778 | 0.6920 | 0.6920 | 0.6920 | 0.6707 |
| $>0.05 - \leq 0.10$ | 0.1036 | 0.0898 | 0.0757 | 0.0757 | 0.0757 | 0.0879 |
| $>0.10 - \leq 0.15$ | 0.0495 | 0.0451 | 0.0423 | 0.0423 | 0.0423 | 0.0457 |
| $>0.15 - \leq 0.20$ | 0.0351 | 0.0277 | 0.0292 | 0.0292 | 0.0292 | 0.0315 |
| $>0.20 - \leq 0.25$ | 0.0242 | 0.0192 | 0.0221 | 0.0221 | 0.0221 | 0.0219 |
| $>0.25 - \leq 0.30$ | 0.0180 | 0.0165 | 0.0171 | 0.0171 | 0.0171 | 0.0174 |
| $>0.30 - \leq 0.35$ | 0.0165 | 0.0138 | 0.0152 | 0.0152 | 0.0152 | 0.0152 |
| $>0.35 - \leq 0.40$ | 0.0120 | 0.0110 | 0.0120 | 0.0120 | 0.0120 | 0.0122 |
| $>0.40 - \leq 0.45$ | 0.0101 | 0.0101 | 0.0103 | 0.0103 | 0.0103 | 0.0102 |
| $>0.45 - \leq 0.50$ | 0.0094 | 0.0092 | 0.0081 | 0.0081 | 0.0081 | 0.0091 |
| $>0.50 - \leq 0.55$ | 0.0076 | 0.0087 | 0.0079 | 0.0079 | 0.0079 | 0.0080 |
| $>0.55 - \leq 0.60$ | 0.0064 | 0.0079 | 0.0064 | 0.0064 | 0.0064 | 0.0070 |
| $>0.60 - \leq 0.65$ | 0.0066 | 0.0076 | 0.0072 | 0.0072 | 0.0072 | 0.0069 |
| $>0.65 - \leq 0.70$ | 0.0067 | 0.0069 | 0.0062 | 0.0062 | 0.0062 | 0.0066 |
| $>0.70 - \leq 0.75$ | 0.0056 | 0.0070 | 0.0066 | 0.0066 | 0.0066 | 0.0062 |
| $>0.75 - \leq 0.80$ | 0.0061 | 0.0062 | 0.0063 | 0.0063 | 0.0063 | 0.0060 |
| $>0.80 - \leq 0.85$ | 0.0060 | 0.0063 | 0.0067 | 0.0067 | 0.0067 | 0.0063 |
| $>0.85 - \leq 0.90$ | 0.0074 | 0.0068 | 0.0065 | 0.0065 | 0.0065 | 0.0068 |
| $>0.90 - \leq 0.95$ | 0.0080 | 0.0073 | 0.0082 | 0.0082 | 0.0082 | 0.0081 |
| $>0.95 - \leq 1.00$ | 0.0180 | 0.0151 | 0.0139 | 0.0139 | 0.0139 | 0.0162 |
| Sum | 1.0000 | 1.0000 | 1.0000 | 1.0000 | 1.0000 | 1.0000 |
| $r^2 > 0.15$ | 0.2040 | 0.1872 | 0.1900 | 0.1886 | 0.2089 | 0.1957 |
| $r^2 > 0.70$ | 0.0512 | 0.0486 | 0.0482 | 0.0486 | 0.0518 | 0.0497 |

### Long-range LD (LRLD)

#### Estimating LRLD by random samples of syntenic marker pairs

Pooled over all families, 418,075 pairs had a distance above 20 Mb; as expected, no high LD of r^2^ ≥ 0.7 was found beyond 20 Mb (Supplemental Table 2).

***Supplemental Table 2.** Distribution of distances among random marker pairs in the F_6_ families*.

Detailed inspection of the distances up to 20 Mb showed that all high LDs were in fact within 10 Mb (Supplemental Table 3).

***Supplemental Table 3.** Distribution of distances among random syntenic pairs with r^2^ ≥ 0.7 separated by up to 20 Mb in the F_6_ families*.

Pooled over all families together, a total of 50,100 random marker pairs qualified within the LRLD definition, namely r^2^ ≥ 0.7 over a distance ≥ 1 Mb. These LRLDs constitute 30.9% of all pairs within 20 Mb, and 1.5% of the total number of syntenic pairs tested (Supplemental Table 2).

Among the syntenic pairs, 0.016 had r^2^ > 0.95, almost 15-times the proportion of the single LD value in this range (0.000001) found among the non-syntenic pairs (Supplemental Table 1). Thus, the proportion of syntenic high LD was not negligible. LRLDs were distributed over all autosomes in all five F_6_ families (Supplemental Table 4, Figure 1; LD matrices in Genetics figshare portal). No LRLD was found on Chromosomes 22 in any of the families.

***Supplemental Table 4.** Distribution of random LRLDs over autosomal chromosomes in the F_6_ families*.

Though these LRLDs were obtained by random sampling of marker pairs, repeated similar locations of marker pairs suggest the existence of many LRLD blocks. This was indeed found by the LD analysis of the MD QTLRs (see below).

#### F_6_ MD QTLRs and random LRLD

To check for a possible relationship between the LRLDs found here and the F_6_ MD QTLRs mapped in the same population (Table 1), LRLDs and QTLRs were aligned together (e.g., Figure 1 and LD matrices in Genetics figshare portal), and overlaps were counted (Supplemental Table 5).

**Figure 1.**
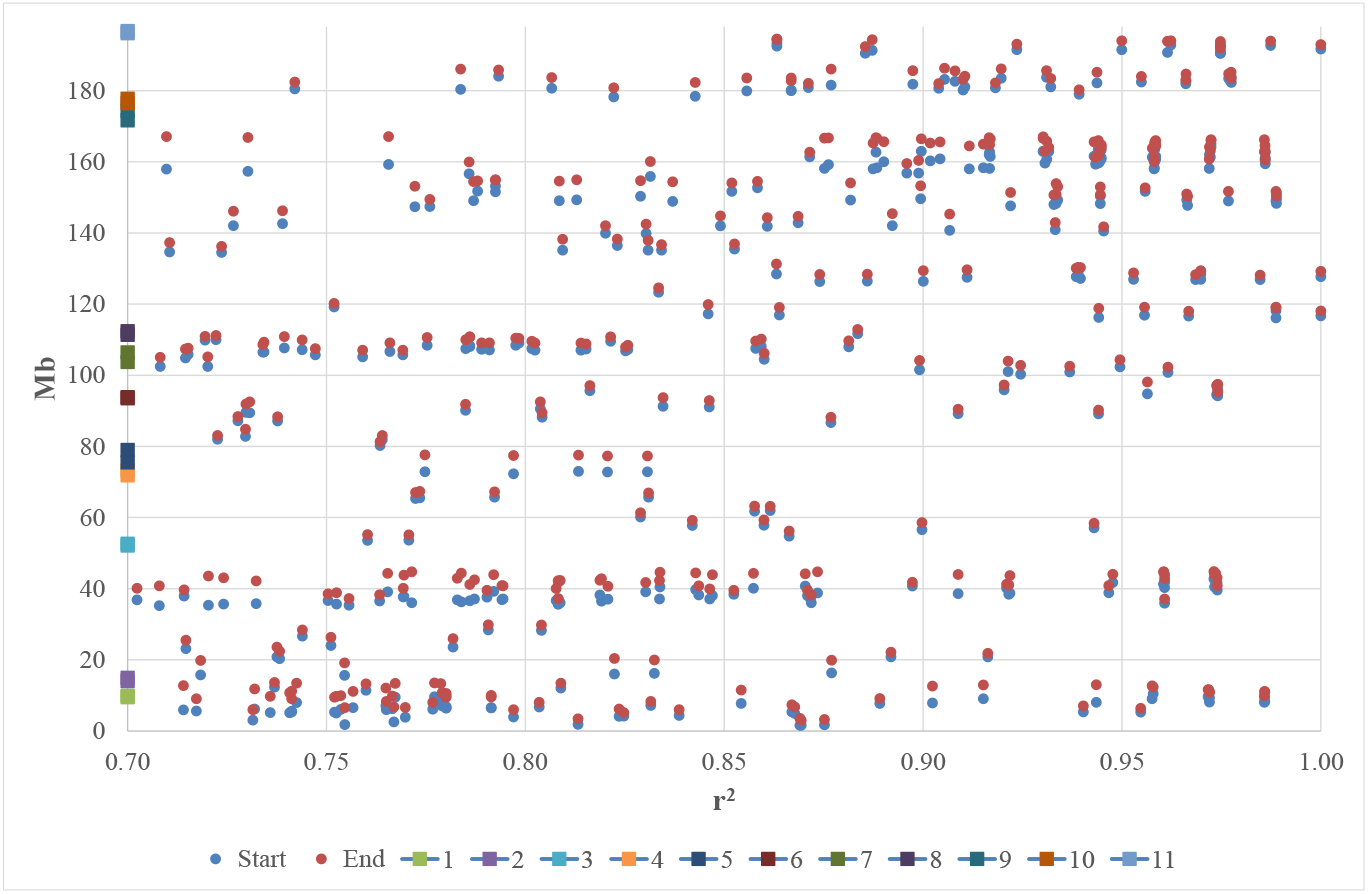
Distribution of random LRLDs over Chr 1 and overlaps with QTLRs (Table 1) in F_6_ Family 1. Location of marker pairs plotted against r^2^. Start, End, locations of the markers in a pair (for each Start dot there is a matched End dot; see Figure 2 for clarity); numbers, QTLR numbers (Table 1).

***Supplemental Table 5.** Number of random F_6_ LRLDs overlapping F_6_ MD QTLRs*.

As noted above, with all markers in an interval less than 1 Mb, no LRLD could be found on Chr 16 (Supplemental Table 4); hence, QTLR 32 was not included in any further analyses. Of the remaining 37 QTLRs, overlaps between 28 QTLRs and LRLDs were found in all families (the non-zeros under ‘Families’ in Supplemental Table 5). It seems remarkable that, even though only 1.5% of the random LD values were LDLR, no less than 75.7% of the mapped MD QTLRs overlapped LRLDs. Then again, in Galgal6, QTLRs averaged 1.4 Mb (Table 1), and random LRLDs averaged 2.2 Mb, from 1 to above 12 Mb (Supplemental Table 6). Thus, such overlap may not be so surprising, but a result of the abundance and size of the QTLRs and LRLDs.

***Supplemental Table 6.** The distribution of random F_6_ LRLDs length*.

Zooming in on QTLRs clearly showed the overlap between the LRLDs and QTLRs (Figure 2). Not only was LRLD found within QTLRs, but LRLD was found between QTLRs 4 and 5 in all 5 families. The similar locations seen in Figure 2 suggest the presence of LD blocks shared by both QTLRs.

**Figure 2.**
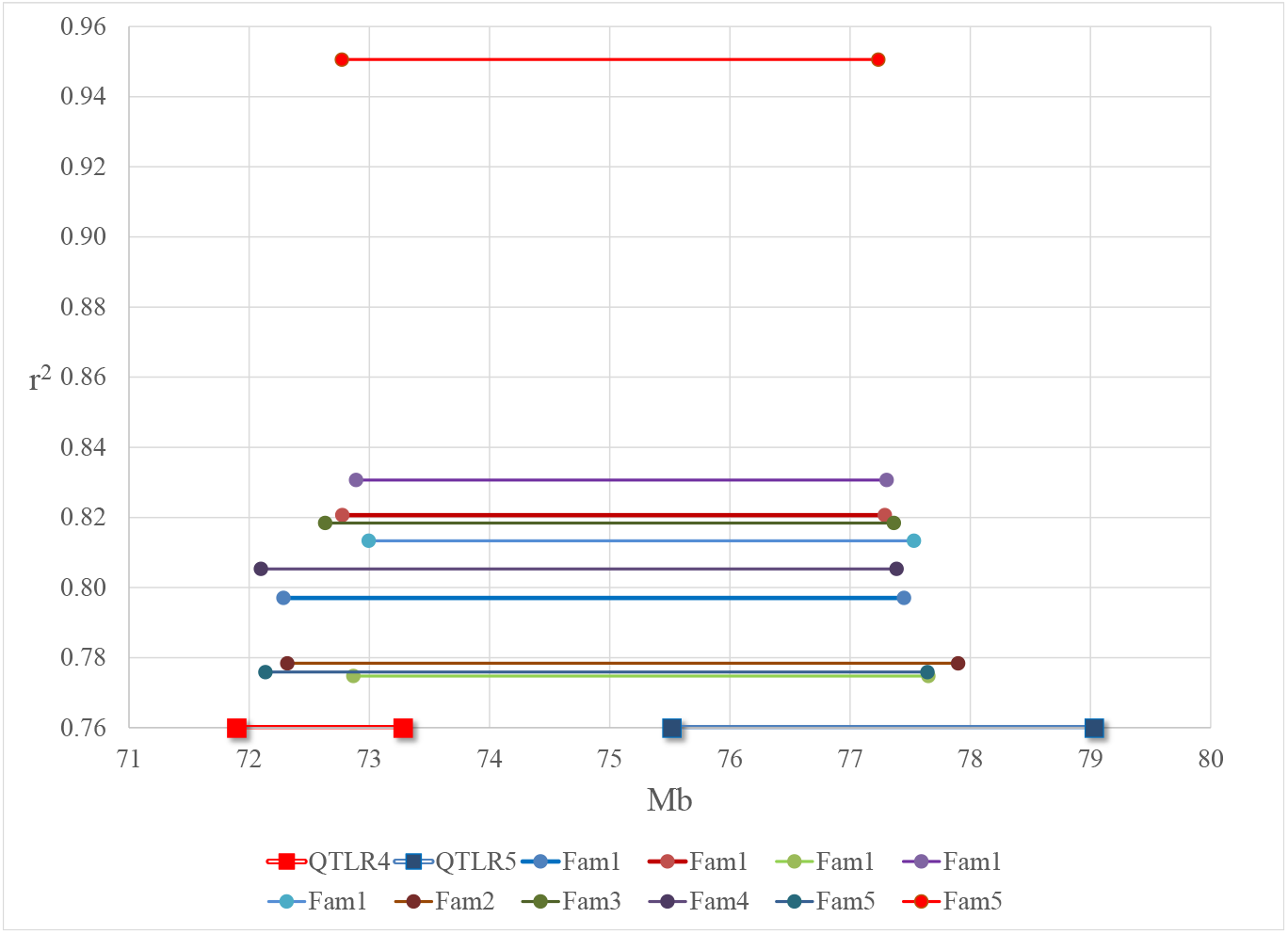
Overlaps between QTLRs 4-5 and LRLDs in all F_6_ families. Fam, family. There were 5 LRLDs in Family 1, 1 LRLD in Families 2 - 4, and 2 LRLDs in Family 5. Each series of an LRLD is composed of two circles connected by a single line; the circles represent the locations on the x-axis of 2 random markers constitute a pair, and the LD r^2^ of this pair is presented on the y-axis. There could be more than one LRLD in a family. Similarly, the series of QTLRs 4 and 5 are presents by squares connected by a double line; the squares represent the locations of the QTLR boundaries. QTLRs do not have an r^2^ of course; they are simply presented on the x-axis, aligned with the LRLD.

### LD in the QTLRs in the F_6_ families

The overlaps found in the F_6_ families between random LRLDs and the MD QTLRs, led us to examine in more detail the LRLD and LD blocks in these QTLRs, with all informative markers of the five F_6_ families (note that this part used *all* pairs of informative markers in the QTLRs, and not only a *sample* of random pairs as in the first LD analysis).

Chromosomes 1, 2, 4, 5, 6 and 14, harbored more than one QTLR (Table 1), thus enabling examination of LD in and between QTLRs. In each F_6_ family, Affymetrix SNP array genotypes were used to calculate LD between all possible pairs of all markers in the 21 QTLRs on those chromosomes.

Hundreds of thousands of LRLDs were found in and between the tested QTLRs (Supplemental Table 7). Total number of marker pairs ranged from below 8 to above 10 million in a family, to a total of more than 43 million pairs. Of these, pooled over all families, 830,182 were LRLDs (62,103 - 227,015 LRLDs in a family). These constitute 0.7 - 2.6% of all pairs in a family, a total of 1.9%, higher than the 1.5% found among the random pairs over all autosomes (Supplemental Table 3).

A total of 161,832 LRLDs were found between QTLRs (Supplemental Table 7), 19.5% of all LRLDs found (0.6 - 24.9 % among the families).

Family 5 is an outlier in Supplemental Table 7, with a much lower number and proportion of total LRLDs and LRLDs across QTLRs compared to the other four families. Further inspection did not identify any source of this difference. Hence, we have no explanation other than sampling variation.

***Supplemental Table 7.** Number of pairs and sum of QTLR LRLDs*.

Pooling all families together, LRLDs were found in all 6 chromosomes examined (Table 5). No LRLD could be found *in* the QTLRs on Chromosomes 5 or 6 (Table 5), as no QTLR there was larger than 1 Mb (Table 1). However, LRLDs *between* QTLRs were also found in those two chromosomes.

**Table 5.** Chromosomes and LRLDs in and between QTLRs. Number of F_6_ array LRLDs in and between QTLRs on the six chromosomes with more than one QTLR. Chr, chromosome; Fam, family; All, All families combined; In QTLRs, LRLDs within the MD QTLRs; Across QTLRs, LRLDs between QTLRs; Total, total number of LRLDs.

| LRLDs | Chr | Fam1 | Fam2 | Fam3 | Fam4 | Fam5 | All |
| --- | --- | --- | --- | --- | --- | --- | --- |
| In QTLRs | 1 | 83,455 | 52,985 | 55,025 | 39,612 | 0 | 231,077 |
|  | 2 | 15,854 | 5,195 | 0 | 0 | 0 | 21,049 |
|  | 4 | 87,859 | 106,654 | 70,245 | 88,091 | 61,605 | 414,454 |
|  | 5 | 0 | 0 | 0 | 0 | 0 | 0 |
|  | 6 | 0 | 0 | 0 | 0 | 0 | 0 |
|  | 14 | 503 | 511 | 489 | 116 | 151 | 1,770 |
|  | Sum | 187,671 | 165,345 | 125,759 | 127,819 | 61,756 | 668,350 |
| Between QTLRs | 1 | 39,244 | 54,674 | 33,243 | 33,937 | 321 | 161,419 |
|  | 2 | 52 | 122 | 1 | 1 | 0 | 176 |
|  | 4 | 26 | 35 | 24 | 33 | 22 | 140 |
|  | 5 | 1 | 1 | 0 | 0 | 0 | 2 |
|  | 6 | 2 | 2 | 2 | 2 | 0 | 8 |
|  | 14 | 19 | 32 | 8 | 24 | 4 | 87 |
|  | Sum | 39,344 | 54,866 | 33,278 | 33,997 | 347 | 161,832 |
| Total | 1 | 122,699 | 107,659 | 88,268 | 73,549 | 321 | 392,496 |
|  | 2 | 15,906 | 5,317 | 1 | 1 | 0 | 21,225 |
|  | 4 | 87,885 | 106,689 | 70,269 | 88,124 | 61,627 | 414,594 |
|  | 5 | 1 | 1 | 0 | 0 | 0 | 2 |
|  | 6 | 2 | 2 | 2 | 2 | 0 | 8 |
|  | 14 | 522 | 543 | 497 | 140 | 155 | 1,857 |
|  | Sum | 227,015 | 220,211 | 159,037 | 161,816 | 62,103 | 830,182 |

In all families, LRLDs were found between most pairs of QTLRs (Supplemental Table 8 a-f). Exceptional among all pairs of QTLRs, an extremely large number of LRLDs (159,413) was found between QTLRs 4 and 5 on Chr 1 in all families, confirming the results of the random samples (Figure 2). The tight LD between these two QTLRs was further confirmed by the LD blocks (below).

***Supplemental Table 8.** Number of arrays F_6_ LRLDs in and between QTLRs in all families*.

Thus, repeating in all F_6_ families, LRLDs were found to be frequent, distributing within and between QTLRs in all chromosomes tested.

### QTLR LD blocks

#### LD Blocks in the F_6_ QTLR

As shown by the data a complicated LD pattern was found in the F_6_ QTLR. Large, fragmented, and interdigitated LD blocks were found in all five families over all six chromosomes examined (LD matrices in Genetics figshare portal). The range and complexity would have been even larger if moderate LD blocks were included, with 0.15 ≤ r^2^ < 0.7.

An example of fragmented interdigitated blocks is presented in Figure 3a. Close examination of the LD found in Family 1 in this region shows the presence of 3 high LD blocks, all fragmented and all interdigitated with one another: Block 1 includes markers with ID numbers 134-141, 143, 145-149, and 151; Block 2 includes markers 142, 144 and 152; Block 3 includes markers 150 and 337. The fact that, despite their apparent fragmentation, these are indeed genuine blocks is shown in Figures 3 c-d. If the markers in Blocks 2 and 3 were not included in the analysis, (e.g., because they were not on the SNP array or were filtered out by the quality control or were not polymorphic in this family), then three clear unambiguous blocks would have been identified.

**Figure 3.**
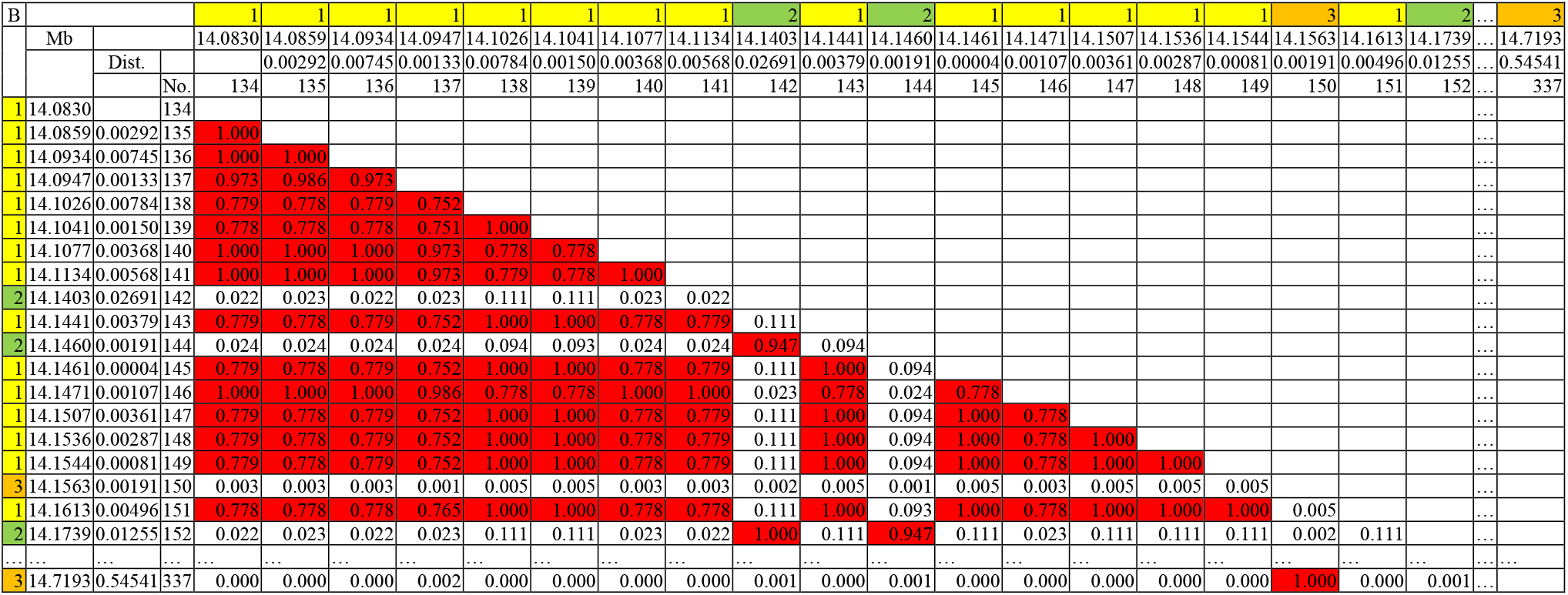
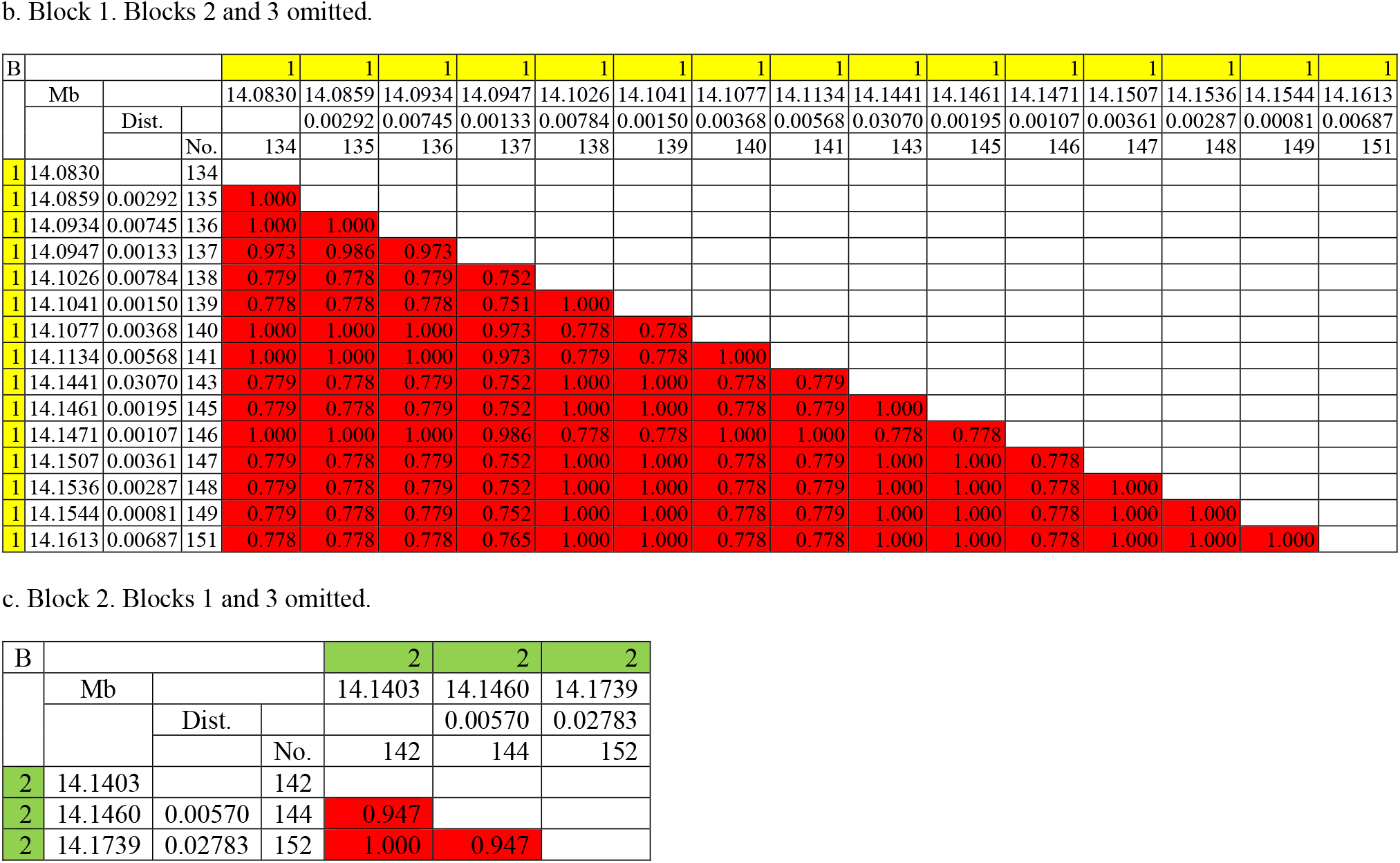
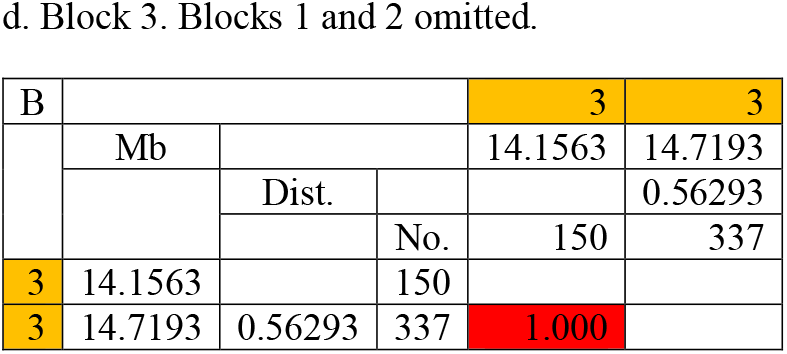
Fragmented interdigitated blocks in QTLR 2 on Chr 1 found in F_6_ Family 1. B, block serial number ordered by the location of the first marker, colored by block; Mb, location on Galgal6 in Mb; Dist., distance in Mb;…, unpresented intermediate markers; No., serial number of the marker; red, LD r^2^ ≥ 0.7.

Note the distance between the markers in block 3, is above 0.5 Mb. Should the criterion of 0.25 Mb [Koch et al., 2013] been used, this block would be defined as LRLD.

#### Blocks shared by QTLRs 4 and 5 in the F_6_ families

In accordance with the random sampling of marker pairs and LRLDs in and between QTLRs in the F_6_, large and long-range LD blocks were shared by QTLRs 4 and 5 in all five families, as exemplified in Supplemental Figure 1 and detailed in Supplemental Tables 8 a-f. In Supplemental Figure 1, the high LD block distributed from the first marker of QTLR 4 to close to the end of QTLR 5, over 5.7 Mb, with 412 markers included. Considering moderate LD of 0.15 ≤ r^2^ < 0.7, would stretch the block all the way to the end of QTLR 5, over more than 7.1 Mb. Thus, the exceptional LD between QTLRs 4 and 5 indicated by the random sample of pairs was confirmed in all F_6_ families by both LRLDs and LD blocks between QTLRs.

***Supplemental Figure 1.** LD blocks shared by QTLRs 4 and 5 in Family 2*.

### LD among QTLR elements in the eight pure lines

LD of elements within and between the F_6_ QTLRs was further examined within eight Hy-Line elite pure lines. Complex LD blocks between elements within and across QTLRs were found, similar to that found in the F_6_ families, over distances from a few bp to a few Mb (Figure 4-6; all LD matrices are in Genetics figshare portal).

**Figure 4.**
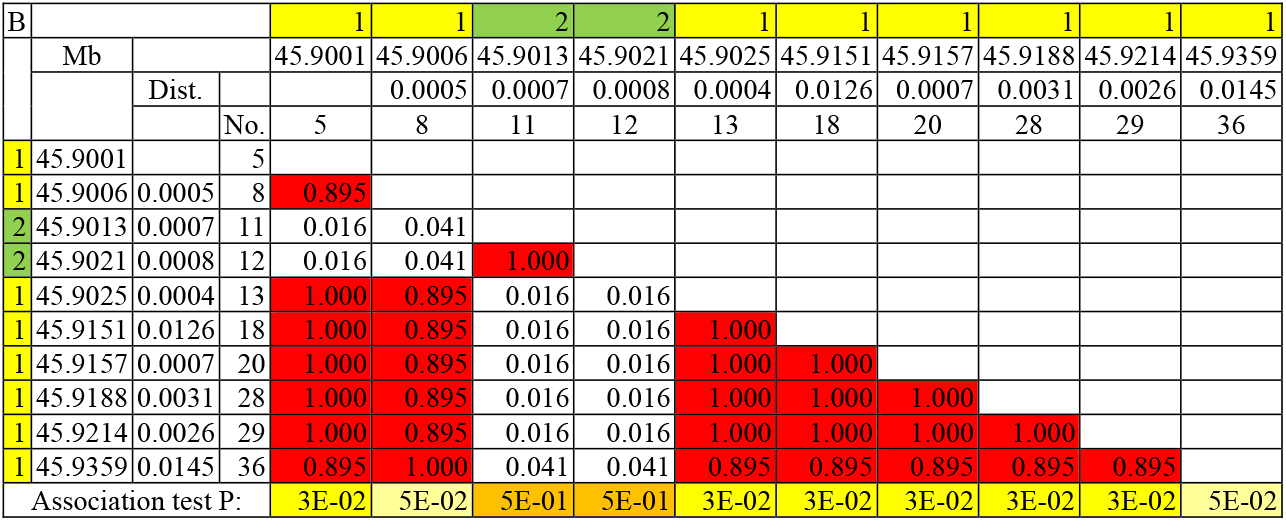
LD within one QTLR gene. Line WL1, the gene TRANK1 in QTLR 13 on chromosome 2. B, block serial number ordered by the location of the first marker (same LD block have the same color); Mb, location on Galgal6 in Mb; Dist., distance in Mb; Red, r^2^ ≥ 0.7; similar P values has similar colors.

**Figure 5.**
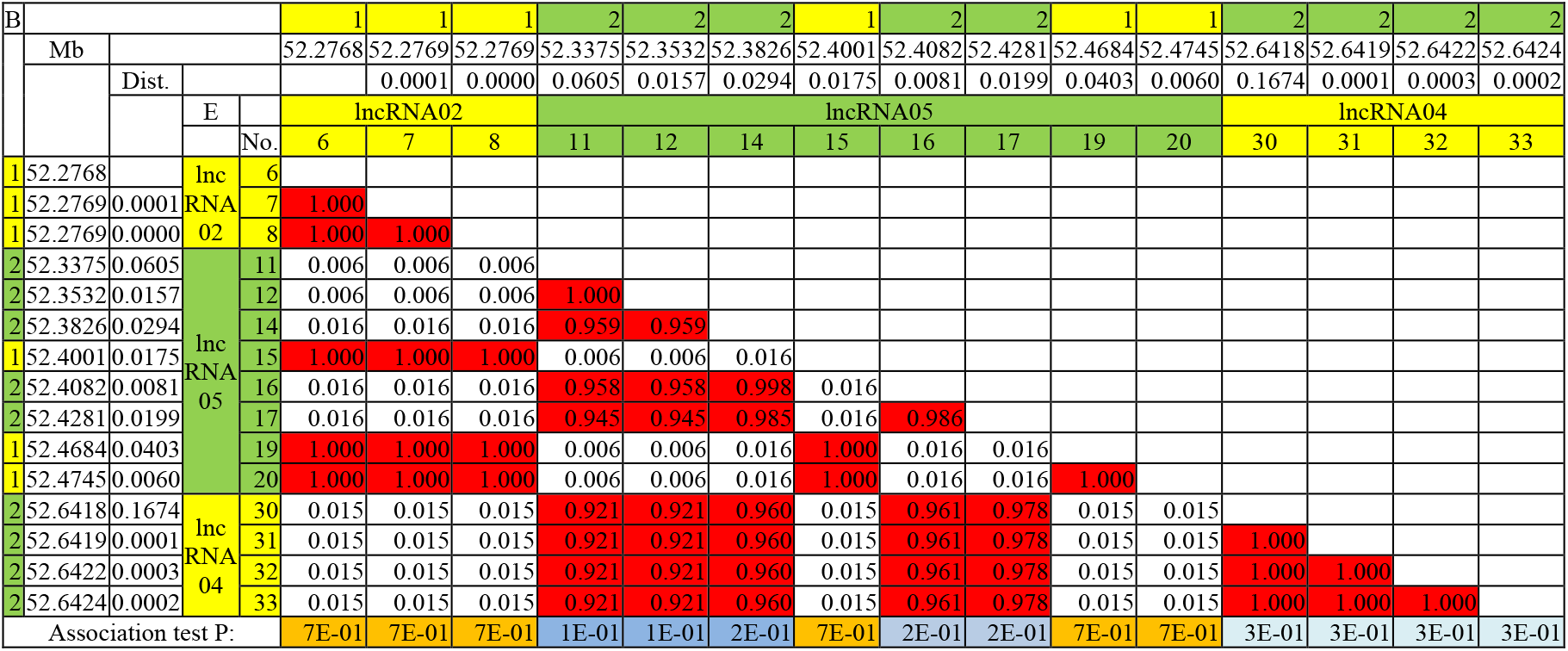
LD blocks across QTLR elements. Line WL3, QTLR 3 on chromosome 1. B, block serial number ordered by the location of the first marker (same LD block have the same color); Mb, location on Galgal6 in Mb; Dist., distance in Mb; E, QTLR element [Smith et al., 2020]; Red, r^2^ ≥ 0.7; similar P values has similar colors.

**Figure 6.**
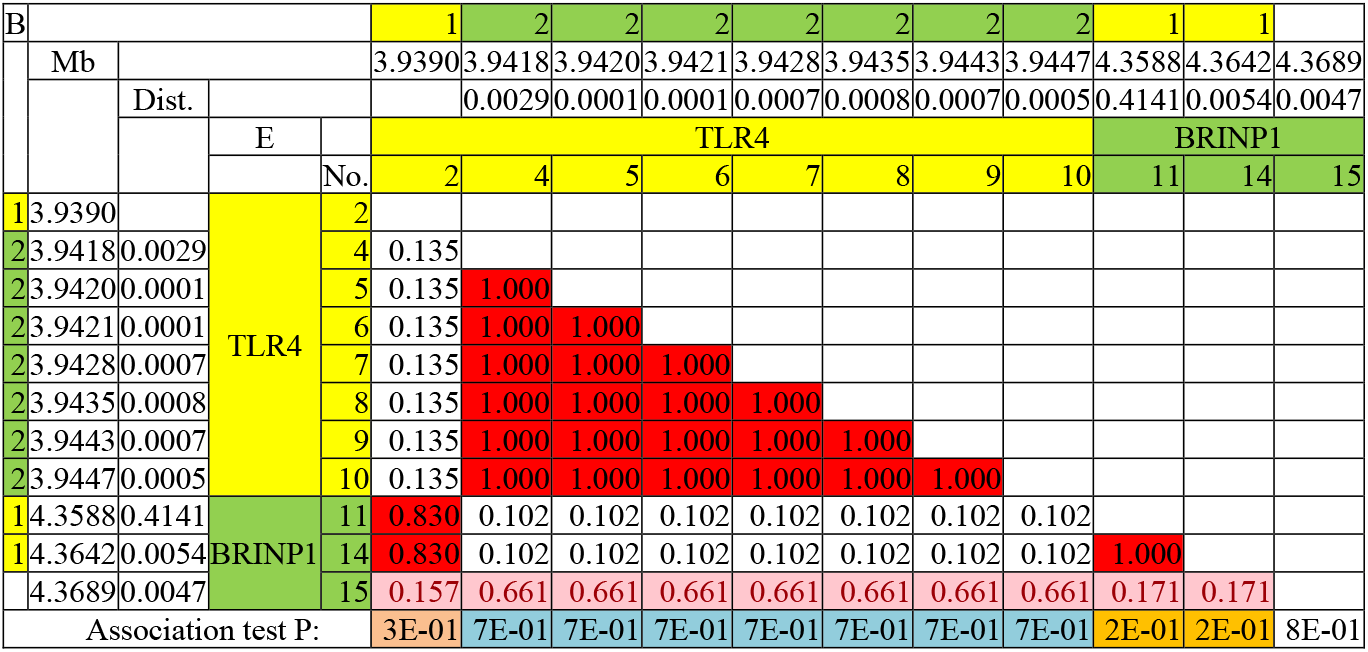
LD between QTLR genes. Line WL2, QTLR 3 on chromosome 17. B, block serial number ordered by the location of the first marker (same LD block have the same color); Mb, location on Galgal6 in Mb; Dist., distance in Mb; E, QTLR element [Smith et al., 2020]; Red, r^2^ ≥ 0.7; purple, 0.15 ≤ r^2^< 0.7; same LD block have the same color; similar P values has similar colors.

#### LD within one QTLR gene

Figure 4 present an example of LD blocks within the QTLR gene *TRANK1* in Line WL1. Despite the short distances (390 bp to 14.5 Kb), a complex pattern was found, with 2 LD blocks, one of which is fragmented around the other. There was high to complete LD between markers 5, 8 and 13-36. These markers had practically no LD with markers 11 and 12, which were in complete LD with one another. Thus, in the gene *TRANK1* in Line WL1, Block 1 starts before, but ends after Block 2. The association test P values [Smith et al., 2020] completely matched the LD blocks, with the same or close P values in each block. This match was found in all other combinations of QTLR - line (Figures 5 - 7).

**Figure 7.**
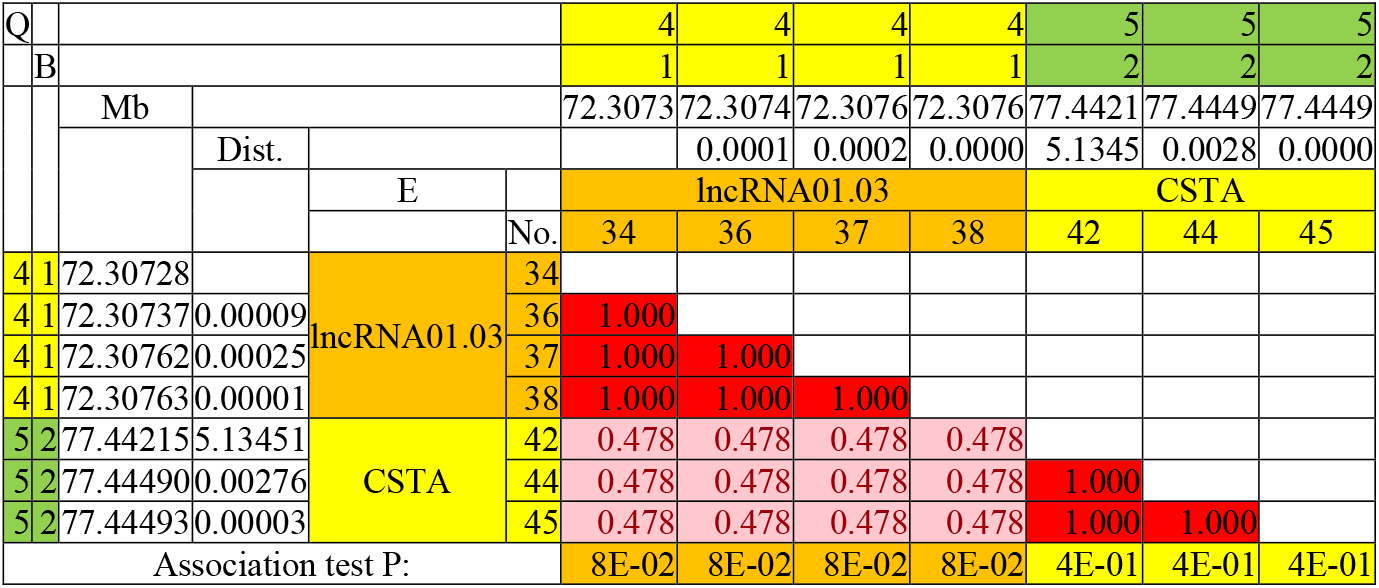
LD between QTLRs 4 and 5. Line WPR1, the lncRNAO1 in QTLR 4 and the gene CSTA in QTLR 5. Q, QTLR serial number (Table 1); B, high LD block serial number ordered by the location of the first marker (same LD block have the same color); Mb, location on Galgal6 in Mb; Dist., distance in Mb; E, QTLR element [Smith et al., 2020]; Red, r^2^ ≥ 0.7; purple, 0.15 ≤ r^2^ < 0.7; same LD block have the same color; similar P values has similar colors.

#### LD between QTLR elements

An example of a more complex LD pattern with interdigitated blocks is shown in Figure 5, this time across QTLR elements (3 lncRNAs).

Careful inspection of Figure 5 shows 2 interdigitated blocks: Block 1 includes Markers 6-8, 15, and 19-20; Block 2 comprise of Markers 11-14, 16-17 and 30-33. Thus, the high LD Block extend over the 3 QTLR lncRNAs. The middle lncRNA05 is split among the 2 blocks. Some of the markers are in LD with upstream lncRNA02, while other markers of the same lncRNA05 form a block with the downstream lncRNA02. The 2 groups of lncRNA05 are interdigitated. That is, Markers 6-8 of lncRNA02 are in the same block with 2 separate regions in the next lncRNA05 - Markers 15 and then 19-20 but not with the other markers in the same lncRNA; Markers 12-14 and 19-20 of lncRNA05 are in LD with all 4 markers of lncRNA04. It would be interesting to find out what are the sources of such complex LD patterns.

LD was found between other types of QTLR elements as well. Figure 6 present such LD between the QTLR genes *TLR4* and *BRINP1* in QTLR 33 on Chr 17. The first marker of *TLR4* has high LD to the first 2 markers of *BRINP1,* and the 3 markers are not linked to other markers of their own gene. The other 6 markers of *TLR4* form a tight LD block. Complexing it even further, the last marker of *BRINP1* (Marker 15) had low to moderate LD with all markers in QTLR 33, both genes included.

#### LD between QTLRs 4 and 5

Markers on both QTLRs 4 and 5 were informative only in Lines WL3, WPR1, WPR2, and RIR1, up to only 4 markers in a line in QTLR 4 (Lines’ LD matrices in Genetics figshare portal). Thus, information on the LD between the QTLRs was limited in this dataset. Nevertheless, in accord with the random LDs in the F_6_ families (Figures 1 and 2) and cross QTLRs LRLD in these families (Supplemental Tables 8 b-f), moderate LD blocks among elements in these QTLRs crossed their boundaries in Lines WPR1, WPR2 and RIR1. In Line WRP1, 2 clear high LD blocks were found, one in each QTLR (Figure 7). However, the 2 QTLR blocks had moderate LDs of r^2^= 0.478 among them, thus forming one moderate LD block. Note that the distances between the cross QTLR pairs, varied from 5.135 to 5.138 Mb.

Looking for a source of such vast, high, and complex long-range LD between QTLRs 4 and 5, a bioinformatics search found 10 and 68 genes in QTLRs 4 and 5, respectively (Figure 8 and Table 6). STRING network analysis revealed five networks of 2 to 28 genes. Two of the networks (‘Net’ 2 and 3 in Table 6), are comprised of genes from both QTLRs (Figure 8 and Table 6). Of ‘Net’ 2, the 2 genes in QTLR 4 and 17 of the 26 genes in QTLR 5 are located in the LD blocks extending over the two QTLRs found in F_6_ (‘+’ in the column ‘B4-5’ in Table 6). Both genes in ‘Net’ 3 are in those blocks. Finally, the two networks with genes from both QTLRs included 6 genes interacting with a gene from another QTLR (Figure 8), all of which located in the cross QTLRs LD blocks. The gene networks and interactions shared by both QTLRs could be the origin of the LD between QTLRs 4 and 5. In fact, the phenomenon of genes whose products work together tending to be on the same chromosomal region is quite common. For example, the Major Histocompatibility Complex (MHC) on chicken chromosome 16 and the Regulators of Complement Activation cluster (RCA) on chromosome 26 [Oshiumi et al, 2005; Hosomichi et al, 2008; Michilak, 2008]. In fact, the networks presented in Figure 8 is a good examples for this colocation of genes working together.

**Table 6.**
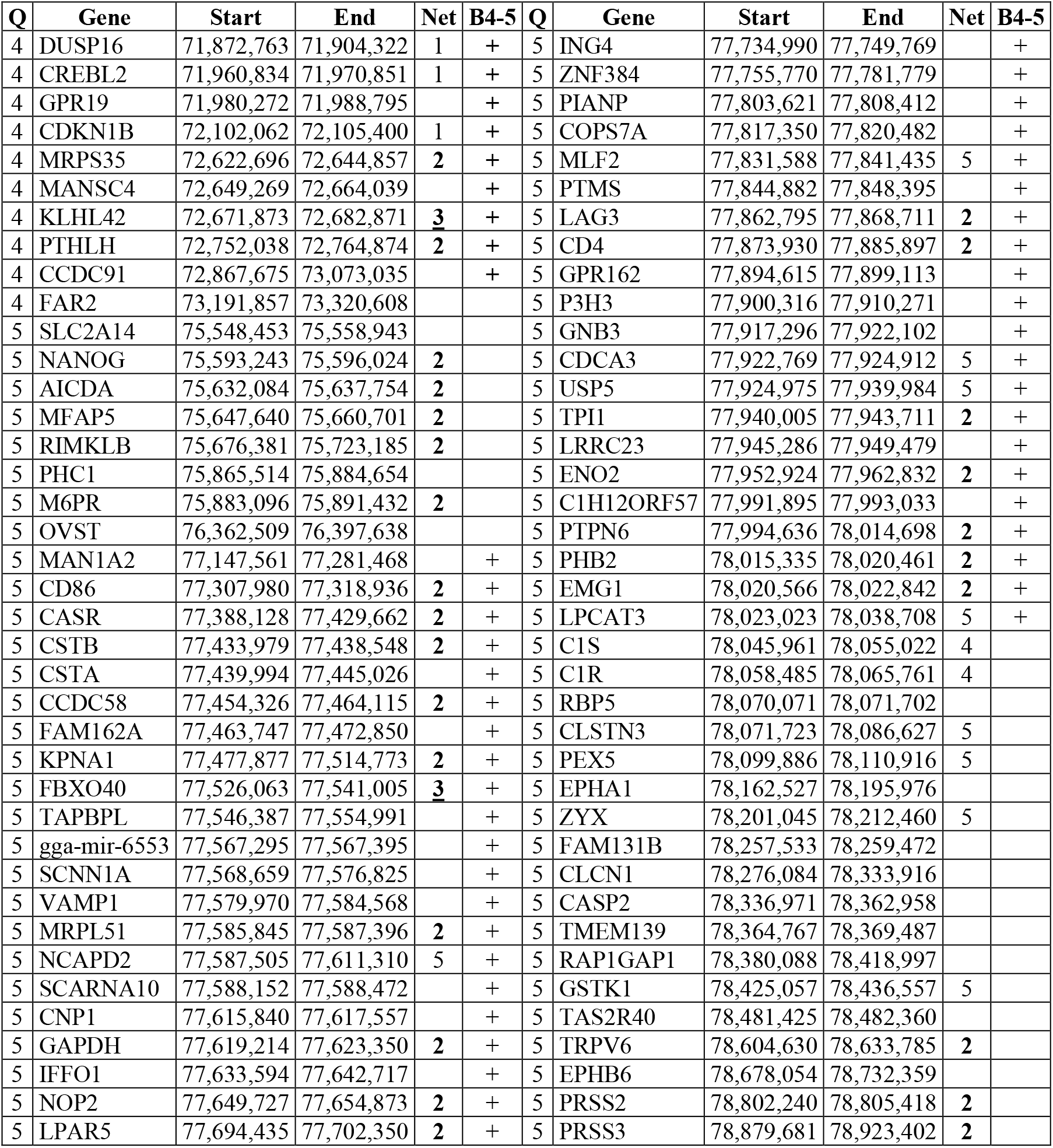
Genes and protein networks in QTLRs 4 and 5, ordered by location. Q, QTLR; Start, End, genes’ coordinated on Galgal6; Net, arbitrary number of a network seen in Figure 8, given by order of location of the first gene (not by order of appearance in Figure 8); Net bolded, net comprise of genes from both QTLRs; B4-5, location in high LD blocks extending over the two QTLRs in the five F_6_ families.

**Figure 8:**
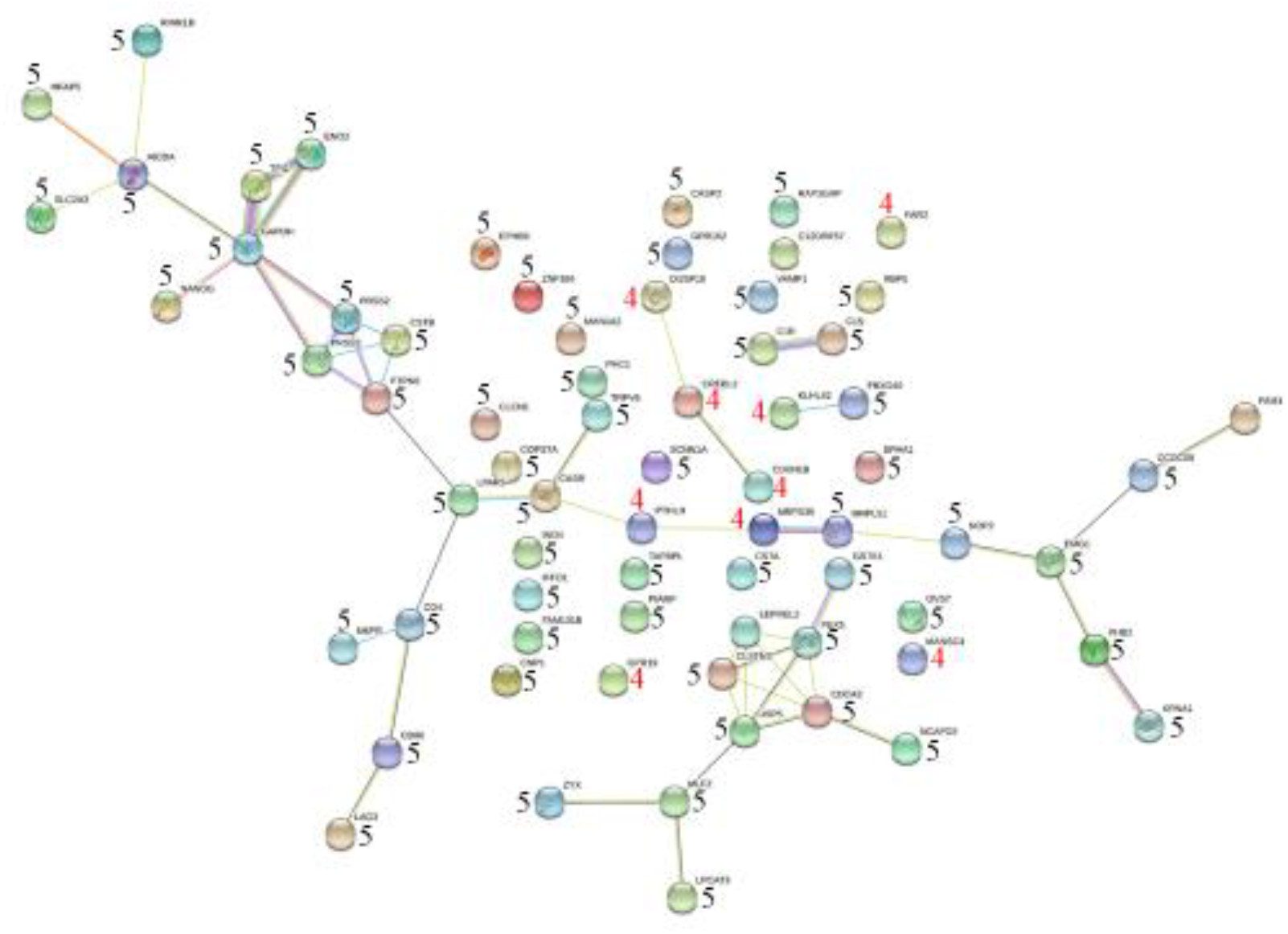
STRING network of genes under QTLRs 4 and 5. Network nodes represent proteins; Colored nodes, query proteins and first shell of interactors; Node content: empty, protein of unknown 3D structure; filled, some 3D structure is known or predicted; Known interactions: 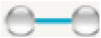; from curated databases; 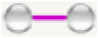, experimentally determined; Predicted interactions: 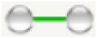, gene neighborhood; 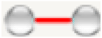, gene fusions; 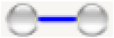, gene co-occurrence; Others: 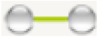, text mining; 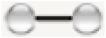, co-expression; 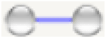, protein homology. The number of the QTLR is shown next to the gene. As can be seen, some gene associations include genes across each QTLR.

## DISCUSSION

Chicken LD over a range of distances, and patterns of LD blocks, were studied in five F_6_ families from a FSAIL, and eight commercial pure layer lines, thus allowing to study the repetition of the results. LRLD was studied in the F_6_ population by random non-syntenic and syntenic samples of marker pairs genotyped by the Affymetrix HD SNP array. In face of the LRLD results, and to illustrate the importance of LRLD to QTL mapping results, LRLD and LD blocks were studied with all possible marker pairs of all markers in previously described MD QTLRs [Smith et al., 2020].

This study started with SNP location information from the previous chicken genome build Galgal4, and was subsequently updated to the Galgal6 assembly. This change necessitated remapping of the QTLRs described in Smith et al. (2020), resulting in negligible changes of most QTLR coordinates. Nevertheless, the change of genome versions moved a segment of Galgal4 QTLR 1 (chr 1) to Galgal6 QTLR 4 (chr 1) (Appendix). The moved segment included one QTLR lncRNA described in Smith et al. (2020), thus emphasizing the importance of basing genomic analyses on the most updated genome version. These results also present the power of LD to identify mapping errors, as already noted by Utsunomiya et al. (2016).

Long-range LD (LRLD) was defined as r^2^ ≥ 0.7 over a distance ≥ 1 Mb. These criteria are more restricted than previously used [Koch et al., 2013; Vallejo et al., 2018]. Nevertheless, repeated appreciable numbers of LRLDs within chromosomes were found repeated in all five F_6_ families by the random sampling of syntenic marker pairs, far above the numbers of high LD found between non-syntenic markers from different chromosomes. The LRLDs were further found in all five F_6_ families by all QTLR array markers on chromosomes with more than one QTLR. These results could be an underestimate, as the F_6_ population was designed to fragment the genome for high-resolution QTL mapping [Heifetz et al. 2007, 2009].

High LD blocks were defined as a group of markers located on the same chromosome, having r^2^ ≥ 0.7 with each other, even if markers with low LD appeared between them. This definition allowed “a look over the horizon” and identification of complex blocks. The phenomenon of fragmented and interdigitated LD blocks were repeatedly found in all five F_6_ families and in all eight pure lines over a vast range of distances, from hundreds of bp to mega bases. The FSAIL population was composed of five families, and they showed similar results. Strength of this analysis was that it repeated five times. Then the same phenomenon was seen in the eight elite lines, further adding strength to the validity of these results, which also agree with previous studies from us and others [e.g., Aerts et al., 2007; Allabi et al., 2005; Lipkin et al., 2013; O’Brien et al., 2014].

A strong linkage was found between QTLRs 4 and 5. LRLD between them was found while analyzing the F_6_ random samples of SNPs within all autosomes. High LD blocks were found in all five F_6_ families, comprised of markers from both QTLRs. Moderate LD blocks between QTLRs 4 and 5 were also found in 3 of the 8 pure lines by QTLR elements’ markers. These results raise the question of what elements on both QTLRs are in high LD over such large distances. Of course, being MD QTLRs, the LD between the QTLRs could be a result of a co-selection for MD resistance. But what make these two QTLRs so different from all other pairs of QTLRs? Why is their LD so exceptional?

To answer this, we looked at the gene content of QTLRs 4 and 5. Ten and 68 genes were found in QTLRs 4 and 5, respectively. STRING protein network analysis revealed five gene networks. Two of the networks include genes from both QTLRs, most of which are located within the LD blocks extending over the two QTLRs found in the F_6_. All 6 genes interacting with a gene from another QTLR are in the cross QTLRs LD blocks. Obviously, the shared gene networks and interactions could be the origin of the LD between QTLRs 4 and 5. However, assessing the uniqueness of the LD between the QTLRs necessitates further study on the distribution of cross QTLR networks and interactions among other pairs of QTLRs with less LD among them. Furthermore, assessing the real effect of the gene networks and interactions needs more molecular, quantitative and population studies. All of these are beyond the scope of the present study.

In general, the complex LD found in this study could stem from technical reasons such as sampling variation. It can also be a result of mapping errors, as was indeed found in this study for regions on chromosome 1 with build 4 (Appendix). However, it could have genuine biological meaning, through processes such as co-evolution of genomic sites (as a result natural or artificial selection), gene conversion, copy number variation, population bottlenecks, non-random mating, and epistasis. The repeatability in different analyses, different datasets and different populations, and the agreement with previous reports strengthen the case for the present results as being a genuine biological phenomenon.

The observation of both LRLD and fragmented interdigitated blocks imply that the causative element is not necessarily the closest, or even close at all to the significant marker, and maybe not even to the significant block of markers. Thus, mapping results and searches for causative elements should be taken with caution, by considering the complexity of the LD. All sites with high LD with a significant marker are in effect candidates to be the causative element.

## Availability of Data

Data have been submitted to the European Nucleotide Archive (ENA) at EMBL-EBI under study accession numbers PRJEB39142 (WGS) and PRJEB39361 (RNAseq). LD matrices are available at figshare.

## APPENDIX Genome assemblies

As stated in the Methods, this analysis started based on the Galgal4 genomic build. Then <u>Lift Genome Annotations</u> tool [Haeussler et al., 2019] of <u>UCSC</u> Browser was used to move from Galgal4 to Galgal6 (GRCg6a). The new coordinates were then used to redefine the F_6_ QTLRs as was done by Smith at al. (2020). The same LD analyses were carried on both assemblies, results were similar, and hence only the results on Galgal6 are presented in the body of the paper. Nevertheless, there was one change worthy of note, and here we show the effect of updating the genome build (moving from Galgal4 to Galgal6), and thus the ability to locate assembly errors by LD.

Using *Galgal4*, almost all high syntenic LDs of r^2^ ≥ 0.7 were below 20 Mb, as expected (Appendix Table 1).

**Appendix Table 1.** Galgal4. Distance distribution of high LD syntenic marker pairs randomized within chromosomes.

| Distance (Mb) | Fam1 | Fam2 | Fam3 | Fam4 | Fam5 | All |
| --- | --- | --- | --- | --- | --- | --- |
| $\geq 20$ | 10,710 | 9,792 | 9,740 | 9,986 | 10,193 | 50,421 |
| >20 - $\geq 30$ | 0 | 0 | 0 | 0 | 0 | 0 |
| >30 - $\geq 40$ | 0 | 0 | 0 | 0 | 0 | 0 |
| >40 - $\geq 50$ | 0 | 0 | 0 | 0 | 0 | 0 |
| >50 - $\geq 60$ | 0 | 0 | 0 | 0 | 0 | 0 |
| **>60 - $\geq 70$** | **1** | **3** | **4** | **3** | **2** | **13** |
| >70 - $\geq 80$ | 0 | 0 | 0 | 0 | 0 | 0 |
| >80 - $\geq 90$ | 0 | 0 | 0 | 0 | 0 | 0 |
| >90 - $\geq 100$ | 0 | 0 | 0 | 0 | 0 | 0 |
| >100 - $\geq 110$ | 0 | 0 | 0 | 0 | 0 | 0 |
| >110 - $\geq 120$ | 0 | 0 | 0 | 0 | 0 | 0 |
| >120 - $\geq 130$ | 0 | 0 | 0 | 0 | 0 | 0 |
| >130 - $\geq 140$ | 0 | 0 | 0 | 0 | 0 | 0 |
| >140 - $\geq 150$ | 0 | 0 | 0 | 0 | 0 | 0 |
| >150 - $\geq 160$ | 0 | 0 | 0 | 0 | 0 | 0 |
| >160 - $\geq 170$ | 0 | 0 | 0 | 0 | 0 | 0 |
| >170 - $\geq 180$ | 0 | 0 | 0 | 0 | 0 | 0 |
| >180 - $\geq 190$ | 0 | 0 | 0 | 0 | 0 | 0 |
| >190 - $\geq 200$ | 0 | 0 | 0 | 0 | 0 | 0 |
| Sum | 10,711 | 9,795 | 9,744 | 9,989 | 10,195 | 50,434 |

Unexpectedly, in all 5 families few high LDs were found with distances from above 60 to 70 Mb (bolded in Appendix Table 1). This was the only interval with high LD above 20 Mb. Detail examination of those 13 extreme long range LDs (LRLDs), showed that all markers in these very distant pairs involved the same 3 regions on chromosome 1. Intriguing, all markers were within or very close to 3 of Smith et al. (2020) F_6_ MD QTL regions (QTLRs). In each pair, one marker was from QTLR 1, the other from QTLRs 4 or 5. Although these 3 QTLRs were significant only in Family 3, the extreme LRLDs were found in all 5 families.

In the face of these results, we LD together all three regions involved (regions of the markers in the 13 LDLRs, not the entire QTLR). As presented in the body of the paper for Galgal6, a very complex LD pattern was obtain in Family 1. A representative example is presented in Appendix Figure 1. The entire matrix of 540 SNPs is too large to present, hence only a sample of markers is presented the give general impression of the LD pattern obtained.

**Appendix Figure 1.**
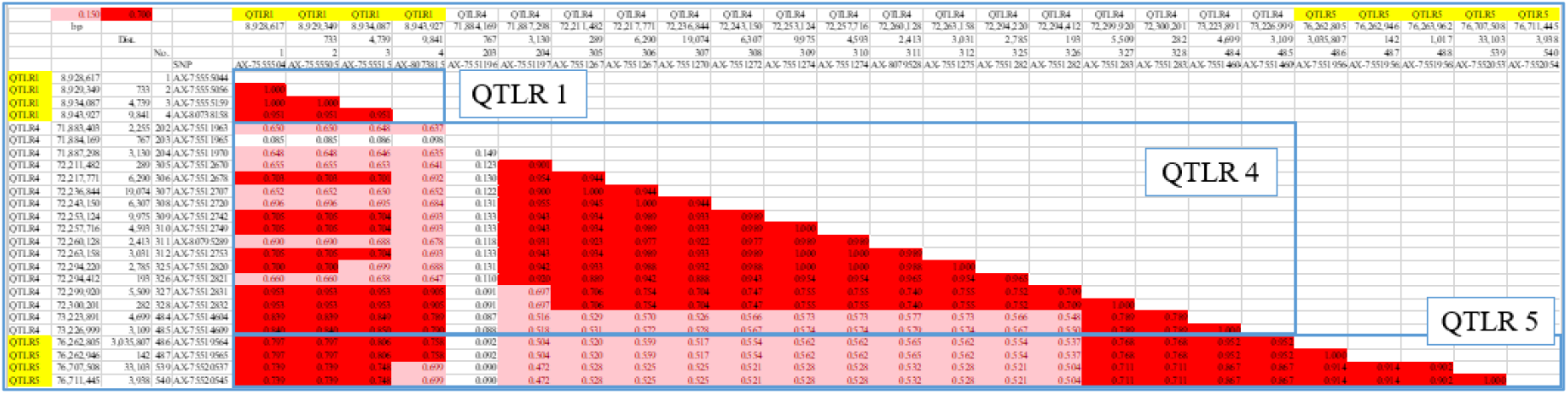
Galgal4. Representative matrix in Family 1 of the 3 regions with markers from LRLDs > 60 Mb. These regions are parts of QTLRs 1, 4 and 5. Yellowed on the flanks, QTLRs 1 and 5, with the white QTLR 4 between them; blue frame, a QTLR; Red, r^2^ ≥ 0.7; purple, 0.15 <r^2^ < 0.7.

On Galgal4 there were over 62.2 Mb between QTLRs 1 and 4, and over 3.0 Mb between 4 and 5. Nevertheless, very high to complete LD values were obtained. As presented in the body of the paper, fragmented interdigitated LD blocks were found, shown by the purple and white rows and columns interfering the red regions.

In light of these results we, LD together the entire 3 QTLRs in each 5 F_6_ family. Appendix Figure 2 present an examples of the analysis of the whole QTLRs, again in Family 1. Similar results with some differences were found in all families.

**Appendix Figure 2.**
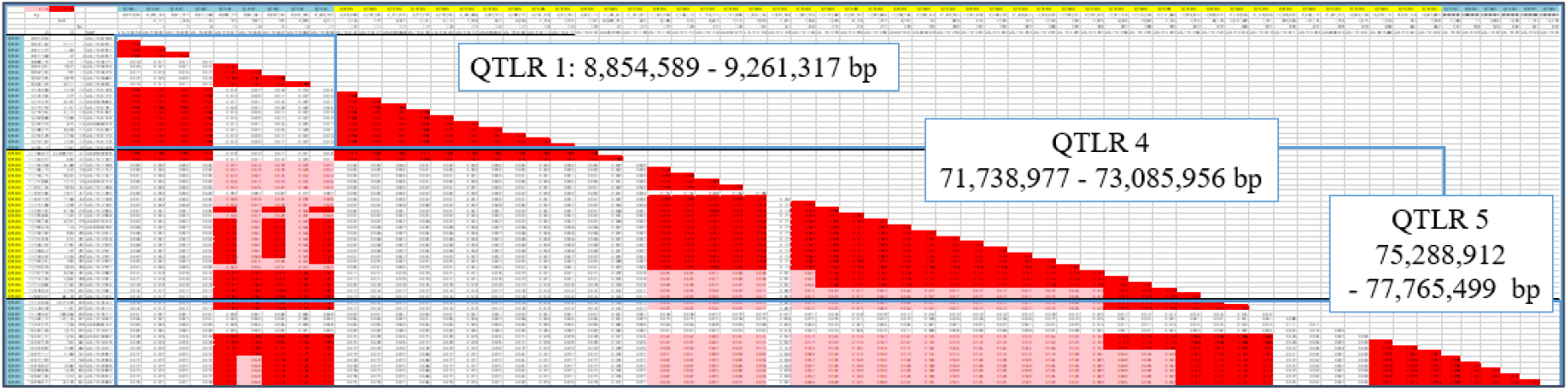
Galgal4. Representative matrix of Family 1 of the entire QTLRs 1, 4 and 5 together. The entire matrix of 602 SNPs is too large to present, hence only a sample of markers is presented to give a panoramic view of the LD pattern obtained. Blued on the flanks, QTLRs 1 and 5 with yellowed QTLR 4 between them; blue frame, QTLR; Text boxes present the boundaries of the QTLRs; Red, LD r^2^ ≥ 0.7; purple, 0.15 <r^2^ < 0.7.

Close examination of Appendix Figure 1 show 2 high LD blocks cover together all QTLR 1, completely unlinked with one another (shown by the white rectangle below and left to the upper left and the lower right red triangles). One of the QTLR 1 blocks is fragmented, covering the start and the end of this QTLR.

The fragmented block of QTLR1 is completely linked with the start of QTLR 4, 62 Mb downstream. The second un-fragmented block of QTLR 1 has moderate to high LD of 0.15 ≥ r^2^ < 0.7 with the fragmented interdigitated blocks of QTLRs 4 and 5.

Moving from Galgal4 to Galgal6, more than 99% of the markers were mapped successfully in Galgal6. Few hundreds markers were deleted in the new assembly, 0.3% of the markers submitted to Lift. Very few markers even changed chromosomes.

After QTLR mapping as in Smith et al. (2020), the same 38 QTLRs reported by Smith el al. (2020) were found. Nevertheless, lnrRNA01 that was located in QTLR 1 on Galgal4 was located in QTLR 4 on Galgal6. Furthermore, Galgal6 completely eliminated the LRLD between QTLR 1 and the other 2 QTLRs, and zeroed that 60 - 70 Mb high LDs (Appendix Table 2).

**Table 4a.** Galgal6. Distance distribution of high LD syntenic marker pairs randomized within chromosomes.

| Distance (Mb) | Fam1 | Fam2 | Fam3 | Fam4 | Fam5 | All |
| --- | --- | --- | --- | --- | --- | --- |
| $\geq 20$ | 10,643 | 9,736 | 9,687 | 9,927 | 10,115 | 50,108 |
| >20 - ≥30 | 0 | 0 | 0 | 0 | 0 | 0 |
| >30 - ≥40 | 0 | 0 | 0 | 0 | 0 | 0 |
| >40 - ≥50 | 0 | 0 | 0 | 0 | 0 | 0 |
| >50 - ≥60 | 0 | 0 | 0 | 0 | 0 | 0 |
| >60 - ≥70 | 0 | 0 | 0 | 0 | 0 | 0 |
| **>70 - ≥80** | **0** | **0** | **0** | **0** | **0** | **0** |
| >80 - ≥90 | 0 | 0 | 0 | 0 | 0 | 0 |
| >90 - ≥100 | 0 | 0 | 0 | 0 | 0 | 0 |
| >100 - ≥110 | 0 | 0 | 0 | 0 | 0 | 0 |
| >110 - ≥120 | 0 | 0 | 0 | 0 | 0 | 0 |
| >120 - ≥130 | 0 | 0 | 0 | 0 | 0 | 0 |
| >130 - ≥140 | 0 | 0 | 0 | 0 | 0 | 0 |
| >140 - ≥150 | 0 | 0 | 0 | 0 | 0 | 0 |
| >150 - ≥160 | 0 | 0 | 0 | 0 | 0 | 0 |
| >160 - ≥170 | 0 | 0 | 0 | 0 | 0 | 0 |
| >170 - ≥180 | 0 | 0 | 0 | 0 | 0 | 0 |
| >180 - ≥190 | 0 | 0 | 0 | 0 | 0 | 0 |
| >190 - ≥200 | 0 | 0 | 0 | 0 | 0 | 0 |
| Sum | 10,643 | 9,736 | 9,687 | 9,927 | 10,115 | 50,108 |

These results show the limitations of any genome build, and present the ability of LD analysis to find assembly errors.

